# Binding language: Structuring sentences through precisely timed oscillatory mechanisms

**DOI:** 10.1101/191494

**Authors:** Katrien Segaert, Ali Mazaheri, Peter Hagoort

**Author notes:** Correspondence to: Katrien Segaert, School of Psychology, University of Birmingham, Edgbaston, Birmingham B15 2TT, UK.

## Abstract

Syntactic binding refers to combining words into larger structures. Using EEG, we investigated the neural processes involved in syntactic binding. Participants were auditorily presented two-word sentences (i.e. a pronoun and pseudoverb such as ‘she dotches’, for which syntactic binding can take place) and wordlists (i.e. two pseudoverbs such as ‘pob dotches’, for which no binding can occur). Comparing these two conditions, we targeted syntactic binding while minimizing contributions of semantic binding and of other cognitive processes such as working memory. We found a converging pattern of results using two distinct analysis approaches: one approach using frequency bands as defined in previous literature, and one data-driven approach in which we looked at the entire range of frequencies between 3-30 Hz without the constraints of pre-defined frequency bands. In the syntactic binding (relative to the wordlist) condition, a power increase was observed in the alpha and beta frequency range shortly preceding the presentation of the target word that requires binding, which was maximal over frontal-central electrodes. Our interpretation is that these signatures reflect that language comprehenders expect the need for binding to occur. Following the presentation of the target word in a syntactic binding context (relative to the wordlist condition), an increase in alpha power maximal over a left lateralized cluster of frontal-temporal electrodes was observed. We suggest that this alpha increase relates to syntactic binding taking place. Taken together, our findings suggest that increases in alpha and beta power are reflections of distinct the neural processes underlying syntactic binding.

## Introduction

For language processing it is common to make a distinction between two crucial components. The first component is memory for the linguistic properties of single words. This knowledge gets encoded and consolidated during language acquisition and is usually referred to as the mental lexicon. Language processing however entails a great deal more than retrieving single words from memory. Rather, the expressive power of human language is based on the ability to combine a limited set of individual words in novel ways to make up new sentences. The binding process that allows us to combine words into larger structures with new and complex meaning, has been referred to as *Merge* (Chomsky, 1995; Grodzinsky & Friederici, 2006) or *Unification* (Hagoort, 2005, 2009, 2016). This binding process is the second crucial component of language processing and is the essence of both language production and comprehension when we use language that goes beyond the single-word level. Binding therefore is a central topic of study in sentence processing research. The process of binding needs to happen at multiple levels. At the level of phonology, words are bound into intonational phrases. Another level of binding is semantic binding: the construction of complex meaning when words are combined into phrases and sentences. A third level is syntactic binding: the combination of words into larger structures, taking into account features that mark syntactic structure, tense, aspect and agreement. Syntactic binding is what we focus on in this paper. Before introducing our study, we would like to highlight the progress that has been made in previous research on the topic of binding.

### The neurobiology of binding

Binding at the most basic level would be the binding of two words in a minimal sentence or phrase. Such minimal binding lies at the core of more complex sentence comprehension and perhaps because it is so fundamental, has attracted increasing research interest in recent years. Bemis and Pylkkänen (2011) investigated the neural changes associated with nouns in a minimal binding context (e.g. red boat) and a wordlist condition (e.g. cup, boat) in an MEG study. Binding for a two-word phrase such as ‘red boat’ involves semantic as well as syntactic binding. For the wordlist condition (i.e. a list of unrelated words), retrieval of lexico-semantic information from memory takes place, but no binding occurs as no phrasal structure can be built for the words in the list. ‘Red boat’ and ‘cup, boat’ thus differ only in the presence versus absence of the binding process: the construction of a syntactic and a semantic relationship. In Bemis and Pylkkänen (2011), wordlists and phrases were presented visually and binding was associated with an evoked response in frontal areas (ventro-medial prefrontal cortex - vmPFC), following an evoked response in temporal areas (left anterior temporal lobe - lATL).

The binding process that takes place when we process a linguistic expression such as a two-word sentence or phrase, is the foundation for processing more complex sentences. The nature of the binding process is invariant. Pushing the complexity down to a minimal two-word paradigm offers the advantage of isolating the binding process from the contribution of other cognitive processes, such as working memory load, which may come into play for longer sentences with complex syntactic structures (see below). The study by Bemis and Pylkkänen (2011) was one of the first studies with a minimal paradigm and inspired many other studies to use a similar design. For example in Bemis and Pylkkänen (2012), binding during both auditory comprehension and reading was associated with an evoked response in the lATL, followed by effects in the left angular gyrus. In this study (unlike in Bemis & Pylkkänen, 2011), no effects were found in vmPFC for the two-word phrase condition compared to the wordlist condition. Pylkkänen, Bemis and Elorrieta (2014) demonstrated that binding in production was associated with parallel effects in lATL and vmPFC. In another key study using a minimal paradigm, using fMRI Zaccarella, Meyer, Makuuchi and Friederici (2015) demonstrated that BA44 is involved in binding not just at the sentence-level, but also at the level of three-word phrases. In all these studies using a minimal paradigm, binding occurred at both the syntactic and the semantic level.

Alongside the research on binding using a minimal paradigm, a lot of studies have been conducted on the neural basis of more complex sentence processing, introducing for example manipulations of syntactic complexity, a priming (repetition suppression) design, or a design with semantic and/or syntactic ambiguities. Mostly, these studies have revealed a left-lateralized network of regions that is associated with the processing of syntactic structures. This network includes primarily the inferior frontal and temporal cortex, and regions in the inferior parietal cortex (e.g. den Ouden et al., 2012; Friederici, Rueschemeyer, Hahne, & Fiebach, 2003; Goucha & Friederici, 2015; Menenti, Gierhan, Segaert, & Hagoort, 2011; Pallier, Devauchelle, & Dehaene, 2011; Papoutsi, Stamatakis, Griffiths, Marslen-Wilson, & Tyler, 2011; Schoot, Menenti, Hagoort, & Segaert, 2014; Segaert, Menenti, Weber, Petersson, & Hagoort, 2012; Shetreet & Friedmann, 2014; Snijders et al., 2009; Tyler, Cheung, Devereux, & Clarke, 2013) (for a review see: Friederici, 2011; Hagoort, 2017; Hagoort & Indefrey, 2014; Tyler & Marslen-Wilson, 2008). In this line of research, some have raised the concern that manipulations of sentence complexity often go hand in hand with variations in working memory load. Indeed, some evidence suggests that activation of the left inferior frontal cortex in particular is related to syntactic working memory rather than syntactic binding (Fiebach, Schlesewsky, & Friederici, 2001; Fiebach, Schlesewsky, Lohmann, von Cramon, & Friederici, 2005). Moreover, others have suggested that the left inferior frontal gyrus is associated with general cognitive functions such as selection among different alternatives, and recovery in the case of misinterpretation (Novick, Trueswell, & Thompson-Schill, 2005; Rodd, Longe, Randall, & Tyler, 2010), rather than with syntactic processing per se. In sum, although some research questions clearly can only be answered using sentences with complex syntactic structures as stimuli, for studies focusing on the process of binding, a minimal sentence paradigm offers clear advantages. A minimal paradigm isolates the binding process from contributions of working memory and other general cognitive functions.

### Isolating syntactic binding using a minimal sentence paradigm

In the present study we use a minimal two-word sentence paradigm to target syntactic binding processes, while minimizing contributions of semantic and phonological binding as much as possible. This is unlike previous studies using a minimal paradigm, which have in common that they all have manipulated binding at multiple levels at the same time. Previous studies have used real words as stimuli. When real words are placed together in a sentence or phrase, both syntactic and semantic binding take place. In this paper, we follow a different approach and use pseudowords. The Dutch ‘tersen’ or English ‘to dotch’ are examples of pseudoverbs. A pseudoword is a unit of text or speech that follows the orthographic and phonological rules of a language, but has no meaning in the mental lexicon. Using a minimal sentence paradigm with pseudowords, we can thus zoom in on binding at the syntactic level. Also, since we use a minimal sentence paradigm, we rule out contributions of general cognitive processes such as working memory.

We will investigate the EEG changes for the following critical comparison: a minimal sentence condition with syntactic binding (e.g. in Dutch: ‘zij terst’, in English: ‘she dotches’), versus a wordlist condition with no syntactic binding (e.g. in Dutch: ‘cil terst’, in English: ‘pob dotches’). Stimuli are presented auditorily. The second word, e.g. ‘terst’, is the target word, the onset of which we timelock our comparison of interest to. Pseudoverbs such as ‘dotches’ (in English) are present in both the sentence and wordlist condition, thus in both conditions morphological parsing occurs of stems and inflectional affixes (i.e. regular inflectional morphology). Inflectional affixes indicate the number and tense for each instance of a pseudoverb, e.g. ‘dotches’ in English is the regular third person singular.

The minimal sentence condition and wordlist condition thus differ from each other only with respect to whether binding takes place. In minimal sentences such as ‘I dotch’, ‘she dotches’ and ‘we dotched’, the presence of a pronoun and the regular inflectional morphology are cues to establish a syntactic structure and for syntactic binding to occur (similar to Goucha & Friederici, 2015). The aspects of syntactic binding we manipulate are: 1) agreement: we establish agreement of number and person between the pronoun and the pseudoverb in the minimal sentence condition, but not in the wordlist condition; and 2) structure building: ‘subject verb’ is a sentence with a syntactic structure, while for wordlists with two verbs we cannot establish a syntactic structure. The paradigm thus allows zooming in on syntactic binding, with only a minimal contribution from semantics (i.e. the pronoun signals who the agent is).

### Oscillitary changes in the EEG related to syntactic processing

We will measure the brain’s response to syntactic binding using EEG. Evoked responses in the EEG (also called event-related potentials, i.e. ERPs) are obtained by averaging event-locked EEG epochs (i.e. trials). An evoked response typically associated with syntactic manipulations is the P600 (Hagoort, Brown, & Groothusen, 1993). Any evoked response reflects the brain’s phase-locked and time-locked response to the experimental event only. A caveat of the event-related averaging approach is that it overlooks activity that is time-locked, yet not phase-locked to the experimental event.

An alternative and complementary way to investigate event-related changes in the EEG signal, is through time-frequency power changes induced in the EEG by the experimental event (i.e. the induced instead of evoked response) (see Bastiaansen, Mazaheri, & Jensen, 2012 for a discussion of these two approaches). Although no study with a minimal paradigm has looked into the oscillatory changes associated with binding, related studies with more complex syntactic manipulations provide insight in the oscillatory changes associated with syntactic processing. Firstly, several studies have suggested that oscillatory changes in the beta range are associated with syntactic binding (see Bastiaansen and Hagoort (2006) and Weiss and Mueller (2012) for reviews on the role of beta in the binding of language). Bastiaansen, Magyari, & Hagoort (2010) found a progressive increase in power in the low-beta band as syntactically correct sentences unfolded. Weiss et al. (2005) found increased low-beta coherence between left-lateralized frontal and temporal regions for the processing of complex versus simpler syntactic structures. Secondly, oscillatory changes in the theta range may play a role. Although changes in this frequency band are more commonly associated with the retrieval of items from the mental lexicon (Bastiaansen, Oostenveld, Jensen, & Hagoort, 2008; Bastiaansen, Van Der Linden, Ter Keurs, Dijkstra, & Hagoort, 2005), Bastiaansen, Magyari, & Hagoort (2010) observed in the theta band (in addition to the beta band) an increase in power as the sentence unfolded, suggesting a role for theta power in binding also. Lastly, both Bastiaansen, Magyari, & Hagoort (2010) and Davidson and Indefrey (2007) observed a reduction in alpha power following a syntactic violation, which could indicate increased syntactic processing after the violation is encountered. Some other studies have related changes in the alpha frequency band to sentence processing: one such study found an increase in alpha band power in auditory sentence processing (Krause et al., 1994), another found increased coherence in the alpha range during sentence reading (Kujala et al., 2007).

In the current study we aim to investigate binding at the syntactic level using a minimal sentence paradigm, to target the precisely timed oscillatory mechanisms associated with syntactic binding. Based on previous literature, we could expect oscillatory changes associated with syntactic binding in the alpha, theta or beta frequency ranges. Given that previous literature does not uniformly point in one direction, we will in our analyses not only be guided by previous research to determine pre-set frequency bands, but also perform a second, complementary analysis that follows a data-driven approach. With a novel utilization of the cluster-randomization test which circumvents multiple comparisons (Maris and Oostenveld, 2007), we examined the power of which frequencies between 3-30 Hz was modulated by syntactic binding. This approach enabled us to assess in which frequencies a significant difference was associated with syntactic binding, eliminating the need to define a-priori frequency bands of interest based on previous literature.

### The present study

In sum, in the present paper, the question is how the brain’s oscillatory changes relate to the syntactic binding process. To investigate the process of binding, we pushed down complexity to the basic two-word sentence level. We compare a minimal sentence condition (e.g. ‘zij terst’, allowing syntactic binding to occur) to a wordlist condition (e.g. ‘cil terst’, no syntactic binding can occur). Through the use of pseudowords, we can zoom in on binding at the syntactic level, minimizing contributions from semantics. The minimal sentence condition and the wordlist condition are constructed as parallel as possible: each form of the pseudoverb is presented in both conditions (i.e. preceded by a pronoun in the minimal sentence condition, preceded by another pseudoverb in the wordlist condition). The critical comparison is for the second word, with the conditions of interest thus only differing in the presence versus absence of the binding process. Our experiment thus allowed investigating the oscillatory mechanisms through which binding of a minimal sentence happens at the syntactic level, largely excluding contributions from binding at other levels, and excluding contributions of other cognitive processes such as working memory.

## Materials and Methods

### Participants

The participants were twenty native Dutch speakers (10 male/10 female, mean age of 21 years with SD 3.1). Participants signed an informed consent that followed the guidelines of the Declaration of Helsinki. The experiments were approved local Ethics Committee of the Social Sciences faculty of the Radboud University (Ethics Approval Number ECG2013-1308-120).

### Materials

Using Wordgen (Duyck, Desmet, Verbeke, & Brysbaert, 2004) we created 20 Dutch pseudoverbs: 'terzen', 'luiven', 'vekken', 'hooven', 'galden', 'gonken', 'golsen', 'zweben', 'zauwen', 'dispen', 'bogsen', 'cillen', 'dunfen', 'ziepen', 'dranen', 'bregen', 'glaven', 'nillen', 'maspen' and 'dernen'. We set Wordgen criteria to generate pseudowords with 6 letters, all ending in -‘en’. All pseudowords could be inflected according to regular inflectional morphology in Dutch, and combined with one of 6 Dutch pronouns, which would yield for example ‘ik ters, ‘jij terst’, ‘hij terst’, ‘zij terst’, ‘wij terzen’, ‘jullie terzen’, ‘zij terzen’.

These materials are inspired by Ullman et al. (1997), who created pseudoverbs in English (e.g. ‘to prass’, ‘to dotch’). Analogous to our Dutch non-verbs, the novel English verbs that Ullman et al. created, could be inflected and combined with pronouns, for example ‘I dotch’, ‘you dotch’ ‘he dotches’, ‘we dotched’.

We made audio recordings of the 6 pronouns and each of the 20 pseudoverbs in the 1^st^, 2^nd^ and 3^rd^ singular and plural present tense. Recordings were made with a female native speaker of Dutch. All created recordings were also saved as a reversed speech version, using the software program Praat (Boersma, 2001) which allows you to play speech in reverse. Again using Praat, we cut pink noise segments matched in length with each individual audio recording of the pronouns and non-words.

### Task and design EEG experiment

Above described stimuli were presented to constitute a minimal sentence condition, a wordlist condition, or one of three filler conditions.

The *minimal sentence condition* consisted of a pronoun paired with a pseudoverb in the 1^st^, 2^nd^ or 3^rd^ singular or plural present tense. The presence of the pronoun and the inflections are cues for syntactic binding to take place (i.e. establishing agreement of number and person between pronoun and pseudoverb, and building of a syntactic structure). In the *wordlist condition* two pseudoverbs were presented, either of which could be in the 1^st^, 2^nd^ or 3^rd^ person. In this case no syntactic binding between the two elements can occur. The EEG experiment consisted of 120 instances of the minimal sentence condition, and 120 instances of the wordlist condition. For the EEG analyses, the analyses contrasts of interest were estimated by comparing the minimal sentence condition and the wordlist condition (i.e. the presence vs. the absence of syntactic binding).

We also used the following filler conditions. A *pink noise condition* consisted of two segments of pink noise which was matched in length with a pronoun and a verb, or with two verbs (120 instances). We included these pink noise trials for variation. We also had a *reversed speech condition* where a segment was played in reverse (90 instances; of these, 30 contained a pronoun and a reversed pseudoverb, 30 contained a reversed pseudoverb followed by a pseudoverb, 15 contained a reversed pseudoverb and pink noise, 15 contained pink noise and a reversed pseudoverb). Lastly, there were 30 instances of a *minimal sentence - mismatch condition*, in which there was no agreement in person and number between the pronoun and pseudoverb. These trials were inserted to ensure some continuity and similarity with the stimuli from the behavioural pretest experiment (see below).

The participants’ task was to detect reversed speech (which only occurred on filler trials). With this task we ensured that participants paid close attention to the stimuli throughout the EEG measurements. Also, there was thus no difference in response decision processes between the crucial conditions of interest, i.e. the minimal sentence and the wordlist condition.

The experimental list consisted of a total of 480 trials, which were preceded by 15 practice trials to gain familiarity with the reversed speech detection task. The presentation time of each element in the trial was as follows (Figure 1): a fixation cross was presented for 200 msec, followed by a grey screen presented for 300 msec, followed by a grey screen presented for 1200 msec of which the onset coincided with auditory presentation of the first word, followed by a grey screen presented for 1400 msec of which the onset coincided with auditory presentation of the second word, followed by a grey screen with two response options (yes/no) presented for 800 msec, followed by a grey screen presented for 1000 msec.

**Figure 1:**
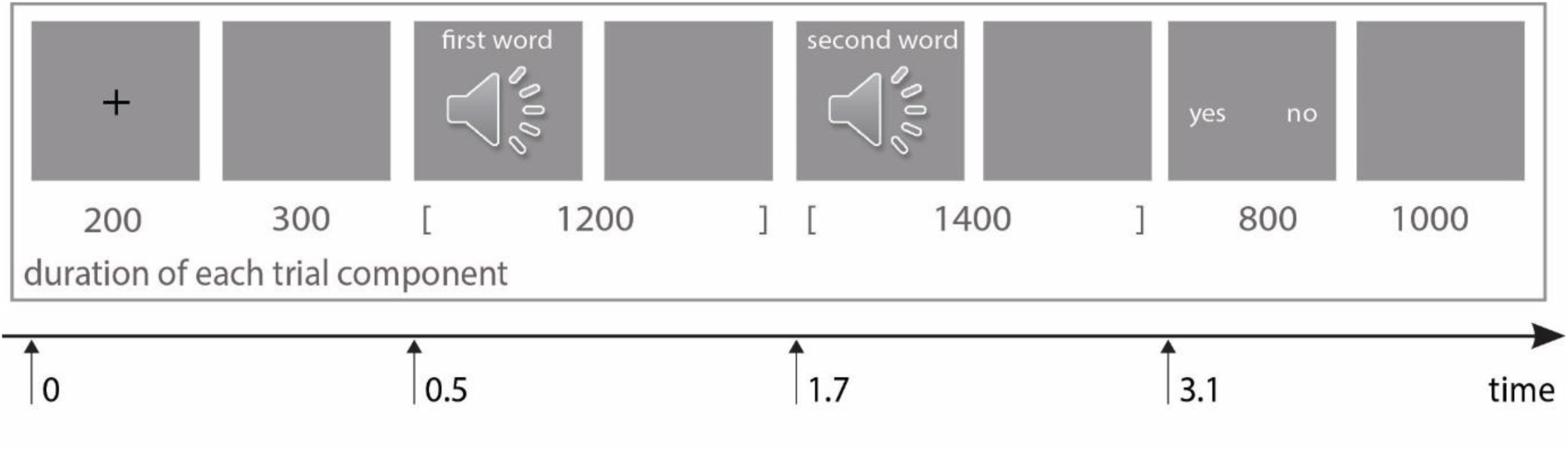
Timing of each component in one trial.

### Behavioural pre-test experiment

Prior to completing the EEG experiment, each participant also completed a short behavioural pre-test experiment. The pre-test experiment was conducted simply to verify that participants indeed are able to syntactically bind minimal sentences containing a pronoun and pseudoverb. This was done by including a minimal sentence condition with agreement mistakes, i.e. *a minimal sentence - mismatch condition*, and instructing participants to detect these mistakes. If participants were able to detect these mistakes, then we could infer from this that participants can perform syntactic binding for a pronoun with a pseudoverb.

The behavioural pretest experiment was made up of three conditions: the minimal sentence condition, the wordlist condition and the minimal sentence - mismatch condition. There were 60 instances of each condition. The experimental list thus consisted of a total of 180 trials, which were preceded by 45 practice trials. The presentation time of each element in the trial was identical in the behavioural pretest and during the EEG experiment (Figure 1).

We will list the results of the pre-test experiment here, since the aim of the pre-test experiment was to validate the manipulation of the main experiment of this paper. The averaged group performance accuracy was 90,0% (SE=1.69%) for detecting agreement mistakes in the minimal sentence –mismatch condition, 92,9% (SE=1.04%) for correctly saying there was no mistake in the minimal sentence condition, and 97,4% (2.24%) for correctly saying there was no mistake in the wordlist condition. Individual performance accuracy was 70% or higher. This suggests that each of the participants was able to perform the task, and thus that each individual participant who completed the EEG experiment, was able to syntactically bind a pronoun with a pseudoverb.

### EEG recording

EEG recordings were made in an electromagnetically shielded room with 60 active surface electrodes placed in an equidistant montage (Acticap, Brain Products, Herrsching, Germany). An electrode on the left mastoid served as a reference; a forehead electrode served as the ground. Vertical and horizontal eye movements were recorded using electrodes on the cap in addition to an electrode placed below the left eye. Using the BrainVision Recorder Professional software (Brain Products GmbH), EEG was sampled at 500 Hz and filtered at 0.2–200 Hz. Impedances were kept below 20 K.

### Data analysis procedure

The offline processing and analyses of the data were performed using functions from EEGLAB version 13.1.1b (Delorme & Makeig, 2004), and the Fieldtrip software package (Oostenveld, Fries, Maris, & Schoffelen, 2011), both are freely available open source Matlab toolboxes. The data was average referenced and epoched from -1.0 s to 5 s after the onset of the visual fixation cue preceding the first word. All trials prior to being sorted into any conditions were visually inspected for non-biological signal artifacts. Ocular and muscle artifacts were removed using independent componentanalysis (infomax algorithm) incorporated as the default “runica” function in EEGLAB 13.1.1b with the first step of a PCA to reduce dimensionality of the data. We have used a similar pipeline for data analysis in previous EEG studies (e.g. van Diepen, Cohen, Denys, & Mazaheri, 2015; van Diepen & Mazaheri, 2017; van Diepen, Miller, Mazaheri, & Geng, 2016).

### Time-frequency representations of power

Using the Fieldrip function ‘ft_freqanalysis_mtmconv’, time-frequency representations (TFRs) of power were calculated for each trial using sliding Hanning tapers having a varying time window of three cycles for each frequency (ΔT = 3/f), an approach which has been used in a number of previous studies (e.g. van Diepen et al., 2015; van Diepen & Mazaheri, 2017; van Diepen et al., 2016; Whitmarsh, Nieuwenhuis, Barendregt, & Jensen, 2011). As to avoid temporal spectral leakage from previous trials, we did not baseline correct the data, but rather compared time-frequency power between the experimental conditions (Mazaheri, Nieuwenhuis, van Dijk, & Jensen, 2009).

### Statistical analysis

We assessed the statistical differences in time-frequency power between the minimal sentence and wordlist condition across participants (random effects analysis) by means of the cluster level randomization test (incorporated in the Fieldtrip software) proposed by Maris and Oostenveld (2007) used in a number of previous studies (e.g. Mazaheri et al., 2009; Nieuwenhuis, Takashima, Oostenveld, Fernández, & Jensen, 2009; van Diepen et al., 2015; van Diepen & Mazaheri, 2017; van Diepen et al., 2016; Van Dijk, Nieuwenhuis, & Jensen, 2010). This is a conservative procedure which circumvents the Type-1 error rate in a situation involving multiple comparisons (i.e. multiple channels and time-frequency points). Here the power of the frequencies of interest, in each channel and time point within the interval 0.5-3.1s after the fixation cross (i.e. the period in between the onset of the first word and the onset of the response screen) was clustered depending on if it exceeded a dependent samples t-test threshold of p<0.05 (two-tailed). Next, the Monte Carlo *p* values of each cluster obtained were calculated on 1,000 random partitions in which the minimal sentence and wordlist condition labels were shuffled.

We followed two analysis approaches. Firstly, we followed an analysis approach in which we were guided by previous literature for the separation in pre-defined frequency bands. We performed analyses within the following frequency bands: theta (4-7Hz), alpha (8-12Hz), low-beta (15-20Hz; Weiss and Mueller (2012) define low-beta as 13-20Hz, we used 15-20Hz to reduce the overlap with the tested alpha band) and high-beta (25-30Hz; Weiss and Mueller (2012) define high-beta as 20-30Hz, we used 25-30Hz to reduce the overlap with the tested low-beta band). In these analyses, we collapsed within the frequency bands to perform cluster level randomization tests.

However, a caveat of collapsing across pre-defined frequency bands is that with such an approach we are possibility overlooking changes in the oscillatory activity that are not falling within the pre-defined selection. As such, in a second analysis approach, rather than collapsing across frequency bands, we assessed statistical differences in power between conditions for each frequency between 3-30 Hz in 1 Hz increments across time and channels. As with the previous approach, power of the frequencies in each channel and time point within the interval 0.5-3.1s after the fixation cross (i.e. the period in between the onset of the first word and the onset of the response screen) was clustered depending on if it exceeded a dependent samples t-test threshold of p<0.05 and the Monte Carlo *p* values of each cluster obtained were calculated on 1,000 random partitions of shuffled conditions.

These two analysis approaches are complementary, as the first is guided by previous research and the second is data-driven. To foreshadow our results, findings from both analysis approaches largely converged.

### Data accessibility

All data and processing scripts will be available upon email request to corresponding author.

## Results

### Behavioral results EEG experiment

During the EEG task, we asked participants to detect reversed speech segments, to ensure they stayed attentive and listened to the stimuli. The averaged group performance accuracy was 95.5% (SE=0.65%) for detecting reversed speech during the filler trials that contained such segments, and 99.6% (SE=0.12%) for correctly answering ‘No’ in all other conditions. Each individual participant performed the task with high accuracy, with the worst scoring participant obtaining 88.9% accuracy for detection of the reversed speech segments.

### EEG results - pre-defined frequency bands

First we will describe the differences in power between the minimal sentence (syntactic binding) and wordlist (no binding) conditions, as revealed by analyses in pre-defined frequency bands as guided by previous literature: theta (4-7Hz), alpha (8-12Hz), low-beta (15-20Hz) and high-beta (25-30Hz). Here we focus first on condition differences *preceding* the onset of the target word (Figure 2), then on condition differences *following* the onset of the target word (Figure 3). The target word is the second word in the two-word sentence, i.e. the word for which syntactic binding occurs.

**Figure 2:**
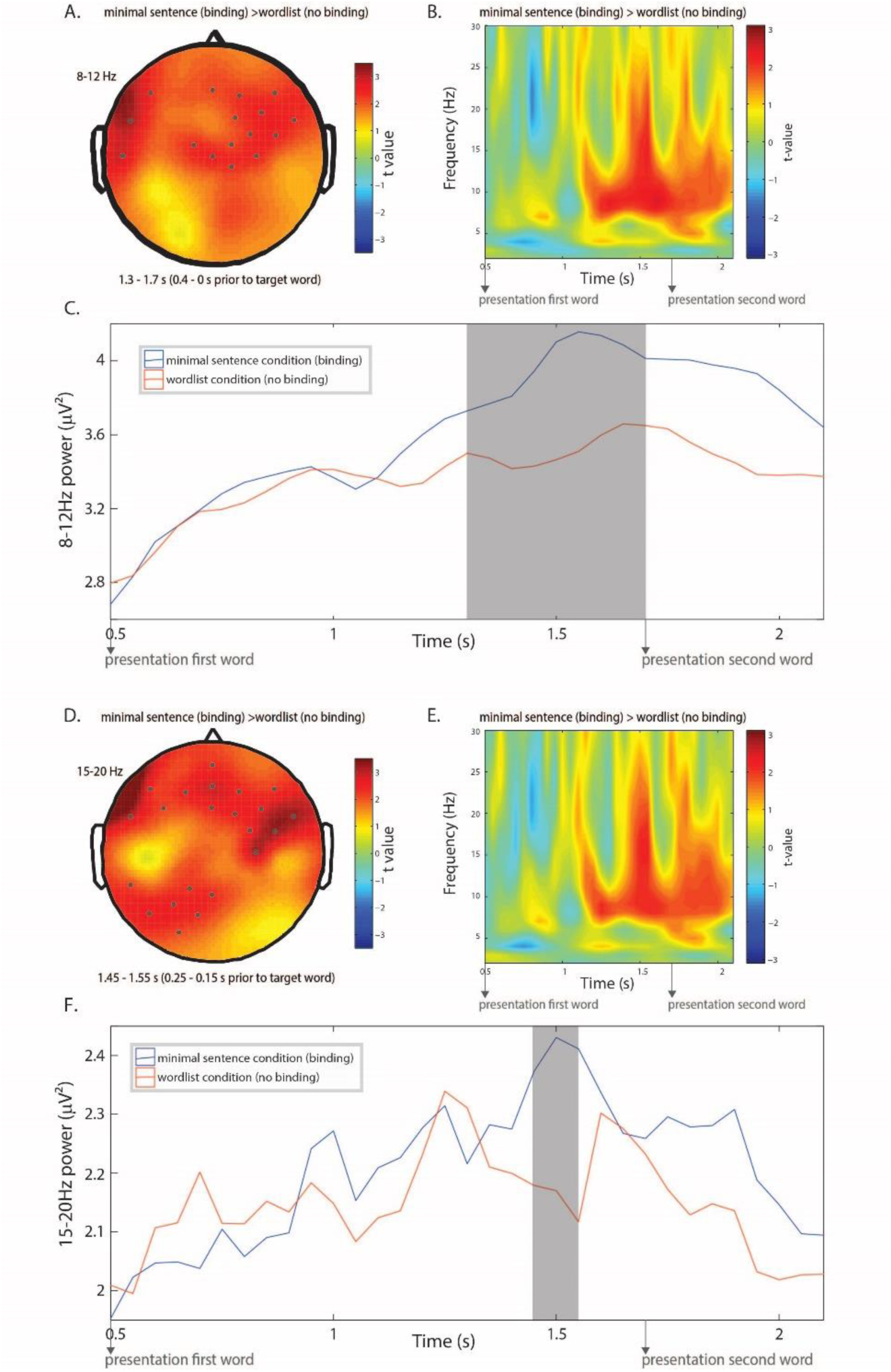
Differences in alpha and low-beta power between the minimal sentence and the wordlist condition *preceding* the onset of the target word. A) For alpha power (8-12 Hz), the scalp topography of the contrast (expressed at t-values) between the minimal sentence and wordlist condition, averaged within the time interval showing a significant power difference (i.e. 0.4 to 0 s prior to the onset of the target word). The dots represent electrodes that showed a significant difference (p<0.05) using the cluster-based permutation test. B) The averaged time-frequency spectrum for the significant electrodes. C) The time course of the power envelope of 8-12 Hz activity (expressed in μV2) of the significant electrodes in both conditions. The grey rectangle indicates the time window in which the difference between conditions is significant (p<.012). D) For low-beta (15-20 Hz) power, the scalp topography of the contrast (expressed at t-values) between the minimal sentence and wordlist condition, averaged within the time interval showing a significant power difference (-0.25 to -0.15 s prior to the onset of the target word). The dots represent electrodes that showed a significant difference (p<0.05) using the cluster-based permutation test. E) The averaged time-frequency spectrum for the significant electrodes. F) The time course of the power envelope of 15-20 HZ activity (expressed in μV2) of the significant electrodes in both conditions. The grey rectangle indicates the time window in which the difference between conditions was significant (p<.039).

**Figure 3:**
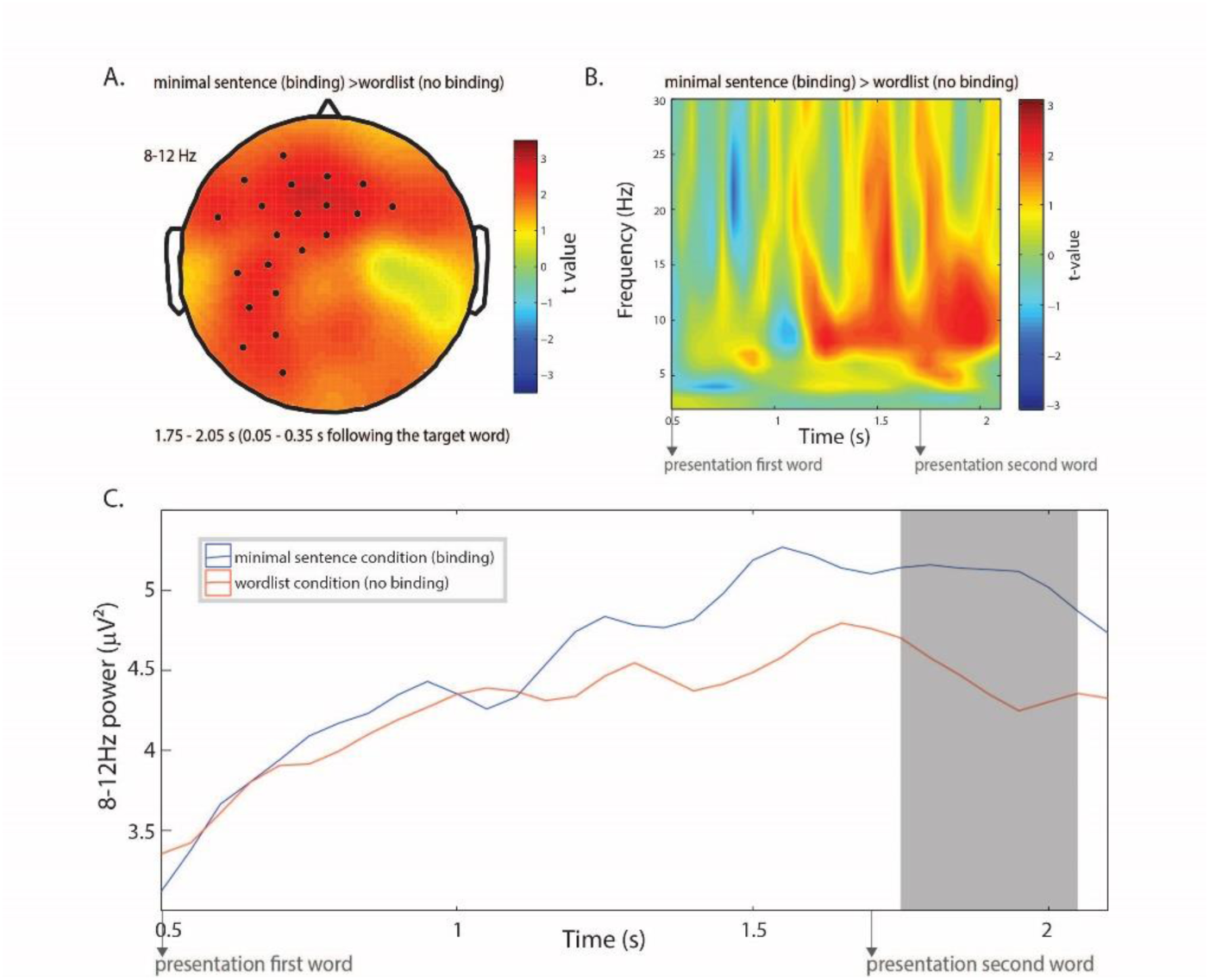
Differences in alpha power between the minimal sentence and the wordlist condition *following* the onset of the target word. A) For alpha power (8-12 Hz), the scalp topography of the contrast (expressed at t-values) between the minimal sentence and wordlist condition, averaged within the time interval showing a significant power difference (0.05 to0.35 s following presentation of the target word). The dots represent electrodes that showed a significant difference (p<0.05) using the cluster-based permutation test. B) The averaged time-frequency spectrum for the significant electrodes. C) The time course of the power envelope of 8-12 Hz activity (expressed in μV2) of the significant electrodes in both conditions. The grey rectangle indicates the time window in which the difference between conditions is significant (p<.035).

*Preceding* the target word, we found no significant difference in the power of *theta (4-7 Hz)* activity between the minimal sentence (syntactic binding) and the wordlist (no binding) condition. We did find a significant difference in the power of *alpha (8-12 Hz)* activity at an interval -0.4 to 0 s preceding the onset of the target word (p<.012) (i.e. time interval 1.3 to 1.7s s from the onset of fixation). Specifically, alpha power was greater in the minimal sentence (syntactic binding) condition, with the difference being most pronounced over a left temporal and central cluster of electrodes (Figure 2A-C). We also found a significant difference in *low-beta (15-20 Hz)* activity at 0.25 to 0.15 s before the onset of the target word (p<.039) (i.e. time interval 1.45 to 1.55s from the onset of fixation). Low-beta power was significantly greater in the minimal sentence (syntactic binding) condition, most pronounced over frontal and parietal electrodes (Figure 2D-F). These analyses revealed no effects in the *high-beta* (25-30 Hz) range.

*Following* the target word, we found no significant difference in the power of *theta (4-7 Hz)* activity between the minimal sentence (syntactic binding) and wordlist (no binding) condition. We did find a significant difference in *alpha (8-12 Hz)* activity at the interval 0.05 to 0.35 s following the target word (p<.035) (i.e. time interval 1.75 to 2.05s from fixation onset). Alpha power was once again larger in the minimal sentence (syntactic binding) condition, this time most pronounced in a left-lateralized frontal-temporal cluster of electrodes. We found no power differences following the target word in *low-beta (15-20 Hz)* or *high-beta (25-30 Hz)* activity between the two conditions.

### EEG results – no pre-defined frequency bands

In addition to the analyses reported above where we collapsed across pre-defined frequency bands as guided by previous literature, we took a complementary approach where we looked at differences between the minimal sentence (syntactic binding) and the wordlist (no binding) condition across every frequency between 3-30 Hz in 1 Hz increments. The spectrogram of Figure 4A illustrates the number of electrodes (at a particular time and frequency) which showed a significant difference in power between the two experimental conditions.

**Figure 4.**
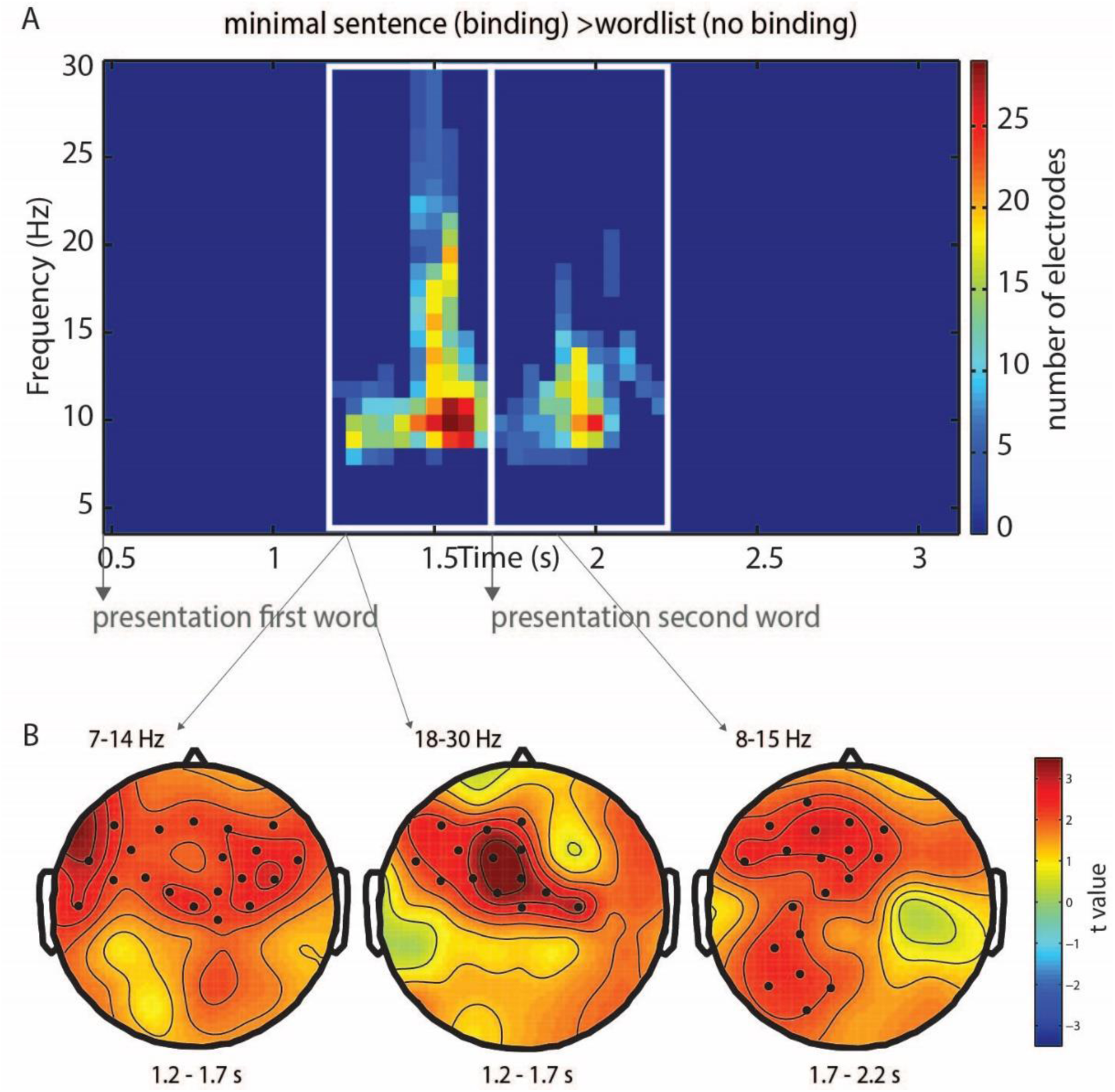
A data-driven approach to assessing the differences in power between the minimal sentence and the wordlist condition. A. Illustrated is the number of electrodes for which a significant difference in power between the minimal sentence (syntactic binding) and the wordlist (no binding) condition was observed, for each frequency and timepoint combination. The values in this graph range from 0 to 26, with the numerical value thus indicating the number of electrodes for which a condition difference was observed for a specific frequency/timepoint. We masked activity that was significant in less than 3 electrodes (value set back to 0). This figure clearly illustrates that preceding the target word, condition differences are observed in the time window from 1.2 s to 1.7 s (left white rectangle). Following the target word, condition differences are observed in the time window from 1.7 to 2.2 s (right white rectangle). B. Averaging over the time window from 1.2 to 1.7 s, significant differences between the minimal sentence and wordlist condition are observed for 7-14 Hz power (condition difference maximal over a central cluster - leftmost topoplot) and for 18-30 Hz power (condition difference maximal over a central cluster - middle topoplot). Averaging over the time window from 1.7 to 2.2 s, significant differences between the minimal sentence and wordlist condition are observed for 8-15 Hz power, most pronounced over left lateralized frontal and temporal electrodes (rightmost topoplot). The dots represent electrodes that showed a significant difference (p < 0.05).

We found that *preceding* the target word, significant differences between the minimal sentence (syntactic binding) and wordlist (no binding) condition were observed in a time window 0.5 s to 0 s before the onset of the target word (i.e. a time window 1.2 s to 1.7 s from the onset of fixation). Averaging over this time window, cluster level randomisation tests revealed significant differences between the minimal sentence and wordlist condition in 7-14 Hz (i.e. alpha) power maximal over a central cluster of electrodes (p<0.043) (Figure 4B, left topoplot). Interestingly, the cluster level randomisation test also revealed a condition difference in the 18-30 Hz (i.e. low-beta and high-beta) power in a central cluster (p<0.024) (Figure 4B, middle topoplot). This suggests that the separation based on pre-defined frequency bands (see above; Figures 2) can sometimes lead to overlooking possible condition effects.

*Following* the onset of the target word, significant differences between the minimal sentence (syntactic binding) and the wordlist (binding) condition were observed in the time window 0 to 0.5 s after the onset of the target word (i.e. a time window 1.7 to 2.2 s from the onset of fixation). Averaging over this time window, cluster level randomisation tests revealed significant condition differences in 8-15 Hz (i.e. alpha) power, maximal over left lateralized frontal-temporal electrodes (p<0.028) (Figure 4B, right topoplot).

In sum, the oscillatory analyses revealed the following: preceding the target word we observed a power increase in the alpha and beta range. Following the target word a power increase in the alpha range is associated with syntactic binding.

### EEG results – P600

Finally, in addition to the oscillatory analyses, we examined any differences in the P600 evoked by presentation of words. Testing for a P600 effect in the latency range from 500 to 700 ms post-word, the cluster-based permutation test did not reveal any significant difference between the minimal sentence and wordlist condition. This suggests that the differences in brain activity we observed above (Figure 2-4) reflect timelocked (to target word onset), but not phase-locked processes.

**Figure 5:**
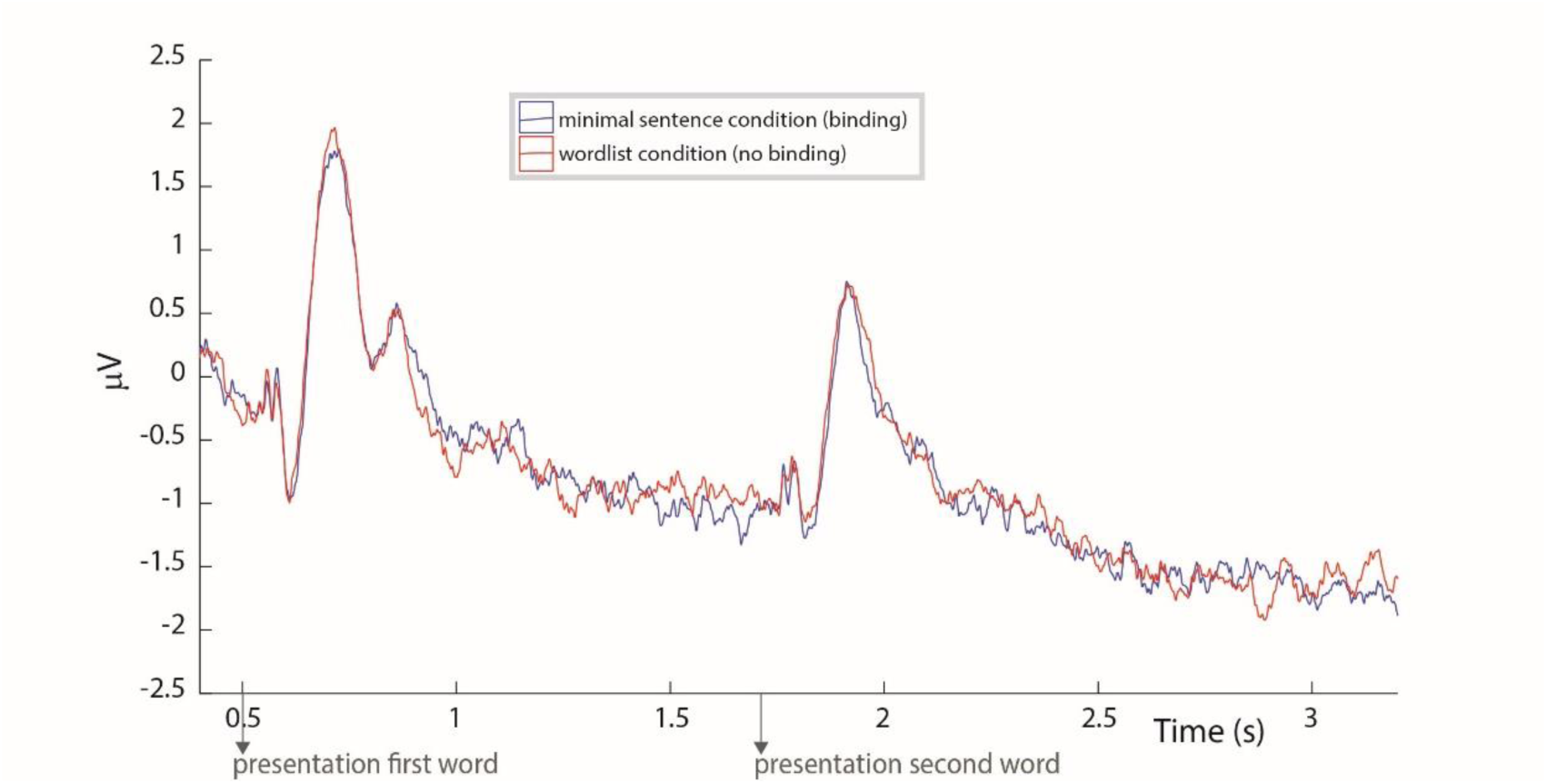
The evoked response in the minimal sentence and the wordlist condition. We did not observe a condition effect on the P600 response. The evoked response is illustrated for two central electrodes (corresponding to electrodes 1 and 2 on the equidistant electrode montage as displayed in Figure 1 of Simanova, van Gerven, Oostenveld and Hagoort (2010)).

## Discussion

In this study we investigated which oscillatory changes in brain activity induced by the onset of words are linked to the syntactic binding process. We focused on the oscillatory changes centered around the second word (target word) in a two-word sentence, comparing the following two critical conditions: a minimal sentence for which syntactic binding occurs (e.g. in Dutch: ‘zij terst’, English equivalent: ‘she dotches’), and a wordlist condition for which no syntactic binding can take place (e.g. in Dutch: ‘cil terst’, English equivalent: ‘pob dotches’). In a minimal sentence such as ‘zij terst’, syntactic binding involves establishing agreement of number and person between pronoun and pseudoverb, and building a syntactic structure. We followed two distinct analysis approaches (one guided by previous literature, and one data-driven approach) and found a largely converging pattern of results. The results can be described as follows. *Preceding* the onset of the target word, in the condition where the target word is to be bound together with the preceding word in the two-word sentence, there was a power increase in the alpha frequency band (8-12 Hz) and low-beta frequency band (15-20 Hz), respectively 0.4-0 s before and 0.25 to 0.15 s before the onset of the target word. Our second analysis confirmed this and revealed that also in the high-beta band there was a significant condition difference preceding the target word (i.e. effects observed 7-14 Hz and 18-30 Hz). The effects preceding the target word were observed mostly in central electrodes. *Following* the onset of the target, there was a power increase in left frontal-temporal electrodes in the alpha frequency band (8-12 Hz) for the minimal sentence compared to the wordlist condition, in a time window 0.05 to 0.35 s following presentation of the target word, signifying the process of binding taking place. Converging findings were observed based on our second analysis approach, i.e. condition difference from 8 to 15 Hz following the target word.

We believe that these findings can be interpreted against the backdrop of theoretical frameworks proposing that a left-lateralized network of frontal-temporal areas is associated with syntactic binding (e.g. Hagoort, 2003, 2009; Tyler & Marslen-Wilson, 2008). Hagoort (2003, 2005, 2009) argues that the left inferior frontal gyrus binds information while the left posterior temporal cortex is responsible for the retrieval of the materials that are needed (e.g. information about syntactic categories, number, gender etc.). Both aspects played a role in our design: for e.g. ‘ters’, information needs to be retrieved to determine that this is the inflection for a first person singular; then, the pseudoverb ‘ters’ is syntactically bound to the preceding pronoun ‘ik’. For target words that could be bound to the preceding pronoun in the minimal two-word sentence (compared to the wordlist condition) we found a power increase in left lateralized frontal-temporal electrodes in the alpha frequency band (8-12 Hz) following the onset of the target word.

We also found increases in power in the alpha, low-beta and high-beta frequency bands immediately preceding the onset of the second word. This could be interpreted as neural responses for the expectation of binding needing to occur (see also Wang, Hagoort and Jensen (under review) for the role of alpha power modulations in predicting upcoming language input). In our paradigm,participants can expect that binding will need to occur when the first word of the two-word sentence is a pronoun. Likewise, when the first word is a pseudoverb, participants can expect that no binding will need to happen. This is analogous to how syntactic binding would occur in natural language processing. When comprehending sentences, words come in one at a time. We expect the sentence to unfold further and can anticipate that upcoming words will need to be bound to the words that are already presented to us. Bastiaansen, Magyari, & Hagoort (2010) observed a progressive increase in power in the theta and low-beta band as the sentence unfolded. We seem to have observed a signature of an expectation for the sentence to unfold further, and binding needing to take place. One interesting implication from the findings of the current study is that future research could explore this signature of expectation further, and investigate whether this relates to for example increased cognitive control, or maintaining information in memory (see also Weiss and Mueller (2012) for a review on different possible functions of beta oscillations during language processing).

The oscillatory mechanisms that emerged in our study are only partly in line with findings of previous studies on syntactic processing in sentences with more complex structures. A common finding in previous literature is an oscillatory change in the beta range associated with syntactic binding (Bastiaansen & Hagoort, 2006; Bastiaansen et al., 2010; Weiss & Mueller, 2012; Weiss et al., 2005). We find an increase in the beta frequency range, not following but directly preceding the word for which binding occurs, which may be a neural signature for the expectation of binding needing to occur. This would be in line with the proposal of a role for beta oscillations in the expectancy of stimuli and maintenance of information (Weiss & Mueller, 2012).

Following the target word, we find an increase in the alpha frequency range associated with syntactic binding. The cluster of electrodes has a left-lateralized frontal-temporal topography, which is in line with several theoretical proposals on binding at the syntactic level (e.g. Hagoort, 2003, 2009; Tyler & Marslen-Wilson, 2008). Weiss and Mueller (2012) proposed a role for beta oscillations in binding, but our findings suggest that this may also extend into the alpha frequency range. Our finding is in line with a previous study showing an alpha power increase for processing auditory sentences (Krause et al., 1994). However, two other previous studies (Bastiaansen et al., 2010; Davidson & Indefrey, 2007) demonstrated that syntactic violations were followed by a reduction in alpha power, suggesting that an alpha suppression effect is associated with increased syntactic processing after the violation is encountered. We however find an increase (not a decrease) in alpha power related to syntactic binding. Our finding can also not readily be reconciled with the commonly attributed role of alpha oscillations to accessing a knowledge system (Klimesch, 2012), since according to this proposal task-relevant regions would show an alpha suppression effect while task-irrelevant regions would show an increase in alpha power. Our findings could suggest that we may be able to attribute an additional mechanism to oscillatory changes in the alpha band, but more research is needed to investigate this.

It must be noted that unlike previous studies investigating oscillatory changes related to syntactic processing, we used a minimal sentence paradigm with pseudoverbs. Such a paradigm offers several advantages. Firstly, with this approach we zoomed in on binding at the syntactic level, with minimal contributions from semantics and phonology. All oscillatory signatures we find can therefore directly be related to the process of syntactic binding. Second, we studied binding at the most basic level and pushed the complexity down to a minimal two-word sentence. There is likely little contribution from general cognitive functions such as working memory (a concern that has been raised for previous studies on syntactic processing using more complex syntactic structures). Future research will have to determine whether our paradigm choice can explain some of the divergence between our findings and the findings of studies investigating oscillatory changes associated with more complex sentence processing.

Though with our paradigm we have zoomed in on syntactic binding, we believe that neither our results nor our interpretation of the results necessarily needs to be specific for syntactic binding only, and not even for the binding of linguistic elements. It may well be that in future studies similar oscillatory mechanisms are observed for semantic, phonological or other forms of binding.

Lastly, we have used two complementary analysis approaches and have showed that both lead to a largely convergent pattern of findings. In our second analysis approach, we were not led by specific frequency bands as defined previously in the literature, but rather looked at each frequency between 3-30 Hz in 1 Hz increments. We uncovered effects in the high-beta range in our second analyses approach, suggesting that a separation based on pre-defined frequency bands could in some cases lead to overlooking condition differences. With this, our paper offers a methodological advance which may be particularly useful in future investigations where researchers do not have a-priori hypotheses about particular frequency bands involved in a task. This may prove particularly useful in a field like psycholinguistics, for which (in comparison to other cognitive domains) relatively little previous literature on neural oscillations is available.

In sum, we investigated the oscillatory mechanisms through which syntactic binding occurs in a minimal sentence paradigm. In the syntactic binding condition, a power increase was observed in the alpha and beta frequency range shortly preceding the presentation of a word that requires binding (relative to when the word cannot be bound to the preceding linguistic context). These signatures may relate to language comprehenders expecting the need for binding to occur. Following the presentation of the target word in a binding context, an increase in alpha power is observed in a left lateralized cluster of frontal-temporal electrodes (a brain network known to be involved in binding). This alpha increase is a neural signature for binding taking place.

## Acknowledgements

Thank you to Laura Arendsen, who created the audio recordings of the stimuli. Thank you to Charlotte Poulisse for assistance in collection of the EEG data.

## Conflict of Interest Statement

There are no conflicts of interest.

## Author contributions

KS and PH designed the study. KS collected the data. KS and AM analyzed the data. KS wrote the manuscript. KS, AM and PH edited the manuscript.

